# Donor embryonic stem cells impede host epiblast specification in 8-cell stage chimeras by crowding and FGF4 signalling

**DOI:** 10.1101/2024.11.08.622647

**Authors:** Stanley E. Strawbridge, Anna Katharina Schrattel, Peter Humphreys, Ken A. Jones, Jérôme Artus, Anna-Katerina Hadjantonakis, Alexander G. Fletcher, Jennifer Nichols

**Affiliations:** Cambridge Stem Cell Institute, University of Cambridge, Jeffrey Cheah Biomedical Centre, Puddicombe Way, Cambridge CB2 0AW, UK; Department of Physiology, Development and Neuroscience, University of Cambridge, Downing Street, Cambridge CB2 3DY, UK; Centre for Stem Cell Biology, School of Bioscience, University of Sheffield, Western Bank, Sheffield, S10 2TN, UK; Department of Cardiology, Heidelberg University, University Hospital Heidelberg, Im Neuenheimer Feld 410, Germany; Cambridge Institute of Science, Altos Labs, Granta Park, Little Abington, Cambridge CB21 6GP, UK; Department of Biochemistry, University of Cambridge, Sanger Building, Tennis Court Road, Cambridge CB2 1GA, UK; Developmental Biology Program, Sloan Kettering Institute, Memorial Sloan Kettering Cancer Center, New York, NY 10065, USA; Université Paris Saclay, Inserm, UMRS1310, 7 rue Guy Moquet, 94800 Villejuif, France; School of Mathematical and Physical Sciences, University of Sheffield, Hicks Building, Hounsfield Road, Sheffield, S3 7RH, UK; MRC Human Genetics Unit, Institute of Genetics and Cancer, University of Edinburgh, Crewe Road, Edinburgh EH4 2XU, UK

**Keywords:** mouse, embryo, trophectoderm, primitive endoderm, mathematical modelling, Bayesian inference

## Abstract

During mouse embryo compaction, outer cells become trophectoderm, while inner cells form the inner cell mass (ICM), later differentiating into primitive endoderm and epiblast - source of embryonic stem cells (ESCs) - during blastocyst formation. Trophectoderm specification is driven by position-governed polarisation, while primitive endoderm specification is positively regulated by FGF4 signalling from the ICM and epiblast. When injected into an 8-cell stage morula, ESCs can exclude host cells from the epiblast, leading to mice derived entirely from these cells. While evidence suggests roles for ESC-produced FGF4 and physical crowding in host cell displacement from the ICM, the interplay between these possible mechanisms has yet to be dissected, in part due to the lack of studies using *Fgf4*^*-/-*^ ESCs. Here, we combine chimaera titration assays with mathematical modelling to study these mechanisms of host cell displacement. Both *Fgf4*^*+/+*^ and *Fgf4*^*-/-*^ ESCs displaced host cells from the epiblast, while only *Fgf4*^*-/-*^ injected embryos reduced primitive endoderm and increased trophectoderm, indicating sequential exclusion by displacement crowding followed by FGF4 signalling.

**SUMMARY STATEMENT:** In mouse blastocyst chimeras, donor cells displace host cells from the epiblast by a combination of crowding and FGF4 signalling.

## INTRODUCTION

The first fate decision in the mouse embryo is driven by positional cues, where outer blastomeres polarise to form the trophectoderm (TE), founding tissue of the placenta (**Fig. 1A, B**). The TE forms an epithelium surrounding the bipotent inner cell mass (ICM) (Hillman et al., 1972; Tarkowski and Wróblewska, 1967), which then segregates into the primitive endoderm (PrE), source of the yolk sac, and epiblast (EPI), future foetus and source of embryonic stem cells (ESCs) (Boroviak and Nichols, 2014; Evans and Kaufman, 1981; Gardner and Rossant, 1979; Martin, 1981). ESCs can be re-incorporated into the pre-implantation embryo by injection or aggregation to form chimeric embryos. These donor cells retain the developmental potential of the pre-implantation EPI and can contribute to all adult germ layers and the germ line. Specification of the ICM occurs asynchronously (Plusa et al., 2008; Saiz et al., 2016), largely due to fibroblast growth factor 4 (FGF4) secretion from unspecified ICM cells and EPI (Feldman et al., 1995; Frankenberg et al., 2011; Kang et al., 2013; Nichols et al., 2009; Yamanaka et al., 2010).

**Figure 1.**
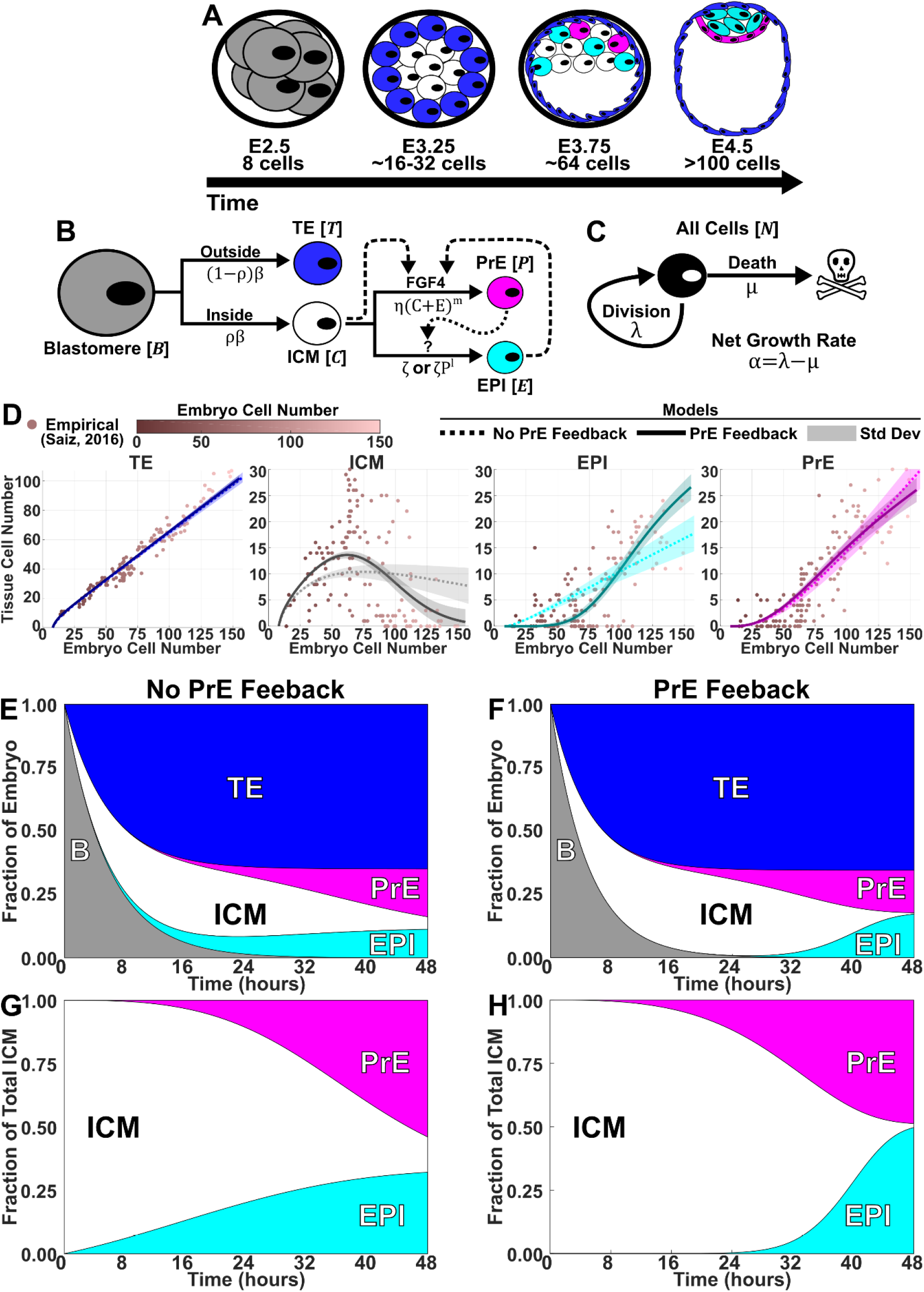
Quantitative modelling reveals a role for feedback from the PrE on ICM specification. **A**. Illustration of pre-implantation mouse development from embryonic day (E)2.5, 8-cell stage morula, to E4.5, late blastocyst. **B**. Model of cell-state transitions during the first two fate decisions during mouse development. Solid lines: cell-state transitions; dashed line: Fgf4 feedback; dotted line: proposed PrE feedback. **C**. All cells grow at net rate α. **D**. Numerical solutions using median posterior parameter values from models without (dotted) / with (solid) PrE feedback, overlaid on data from (Saiz et al., 2016). **E-H**. Reconstructed timecourse of relative tissue sizes in whole embryo (**E, F**) and ICM (**G, H**) for models without (**E, G**) / with (**F, H**) PrE feedback.

The dynamic allocation of the three lineages can be perturbed in myriad ways. Injecting donor ESCs into 8-cell embryos at embryonic day (E) 2.5 can increase TE cell numbers by driving more host blastomeres to an outer position (**Fig. 1B**) (Humięcka et al., 2016). Additionally, the donor cells can increase host-derived PrE cell numbers and host-derived EPI cell numbers. If enough donor cells are injected, the resulting mouse can be composed entirely of donor cells (Poueymirou et al., 2007). This modulation in the second fate decision likely stems from FGF4 production by donor ESCs. Indeed, high, non-physiological concentrations of exogenous FGF4 can drive the entire specifying ICM to the PrE fate (Yamanaka et al., 2010). While FGF4 signalling is known to be a main driver of ICM specification, no feedback role has been proposed for PrE in ICM specification. However, laser ablation studies show that the specifying ICM will compensate in response to loss of EPI or PrE (Saiz et al., 2020), suggesting some form of population-level feedback between the three cell types within the ICM.

Here we unite these observations into a theoretical framework. We first generated a compartment model of cell population dynamics in the mouse embryo from E2.5 to E4.5, calibrated against previous observations (Saiz et al., 2016), indicating a role for PrE feedback on ICM specification. Building on this model, we conducted donor cell injections into host embryos using wild type (WT) *Fgf4*^*+/+*^ and *Fgf4*^*-/-*^ ESCs. Both *Fgf4*^*+/+*^ and *Fgf4*^*-/-*^ donor ESCs can impede host cells from contributing to the epiblast, with *Fgf4*^*-/-*^ donor ESCs yielding a smaller PrE and larger TE. Finally, we combine our base model and chimera assays to generate a model of chimera formation, suggesting that donor cells perturb host cell allocation through spatial crowding followed by FGF4 induction.

## RESULTS AND DISCUSSION

### A compartment model of blastocyst generation suggests a role for feedback from the PrE on ICM specification

Previous models of blastocyst generation have focused on position for the first fate-decision and a bistable gene regulatory network for ICM specification (Bessonnard et al., 2014; Nissen et al., 2017; Saiz et al., 2020). Here we aim to create a minimal model of blastocyst formation, extendable to generate *in silico* chimeras by adding donor ESCs. This model provides outputs for transition rates and numbers and proportions of cells in each lineage. We design a compartment model of mouse embryogenesis spanning the E2.5 8-cell stage to the E4.5 late blastocyst stage (**Fig. 1B**). During this period, two binary cell-fate decisions occur. First, blastomeres, *B*, specify, at a rate **β**, into unspecified ICM cells, *C*, with a bias of **ρ**, or TE, *T*, with a bias of 1 − **ρ**. Second, unspecified ICM cells become either PrE, *P*, or EPI, *E*. PrE specification is driven by FGF4 secreted from the ICM and EPI. This is reflected in the PrE specification rate as η (*C* + *E*)^*m*^, where η is a constant and *m* is a feedback parameter to allow for potential non-linearity. For EPI specification, we compare two models. The first is a constant rate of specification, ζ. This emulates one school of thought that the ICM will take on the EPI identity by default in the absence of FGF4. The alternative model we propose is a yet to be determined feedback from the PrE, possibly ECM components like laminin. In this model the specification rate is ζ *P*^*l*^, where *l* is a feedback parameter to allow for potential non-linearity. Finally, all cells proliferate with the same net growth rate **α** (**Fig. 1C**).

We infer parameters for the two models using an Approximate Bayesian Computation method based on Markov Chain Monte Carlo (ABC MCMC) and data from (Saiz et al., 2016) (**Fig. 1D**). The growth rates of both models were around 0.06 h^-1^ which yields a reasonable doubling time of about 11 h. With this doubling rate the 8-cell morula would double roughly 4 times to yield an embryo with approximately 124 cells. Furthermore, both models capture the fraction of TE (65%) to cells in the ICM compartment (35%), with an ICM bias **ρ** = 0. 35. Feedback parameters were allowed to vary between 0, 1, and 2. In all instances, the parameter inference algorithm selected the feedback parameter to be nonlinear with a value of 2. Both models recapitulate the TE and PrE relatively well (**Fig. 1E-H**). However, the model without PrE feedback does not capture the dynamics of the ICM and EPI. This is especially pronounced when considering unspecified ICM, because by E4.5, most, if not all, cells in the ICM should be either EPI or PrE (**Fig. 1D,G,H**).

This suggests that PrE feedback is necessary to drive timely ICM specification. We aimed to expand on this base model to understand how donor cells perturb host cell allocation. To do this we generated two data sets exploring the host-donor relationship and the modulation of different tissues in response to donor cells.

### Both *Fgf4*^*+/+*^ and *Fgf4*^*-/-*^ donor ESCs can impede host cells from contributing to the EPI

Previous work has shown that donor ESCs have the ability to displace host cells from epiblast through FGF4 signalling ((Poueymirou et al., 2007), (Humięcka et al., 2016)). To disentangle FGF4 signalling from any other mechanism, we investigated host epiblast displacement by injecting either 10 *Fgf4*^*+/+*^ or 10 *Fgf4*^*-/-*^ ESCs into wild-type (WT) 8-cell stage morulae, with intact zona pellucida (**Fig. 2A**). Chimeric embryos were cultured *ex utero*, alongside WT non-injected embryos, for 48 hours to the late blastocyst stage. Embryos were then fixed, immunostained, and imaged for EPI marker SOX2, PrE marker GATA4, and donor cell marker DsRed (**Fig. 2B**). ICM cell numbers and types were determined manually with ImageJ plugin “Cell Counter” (**Fig. 2C, D**). Two groups with varying levels of chimerism were observed in the 10 *Fgf4*^*-/-*^ ESC injected (10-/-) embryos. This observation was confirmed by k-means clustering optimization over the silhouette index (**Fig. S2**). The two resulting groups were chimeras with high (Hi) and low (Lo) levels of donor cell contribution to the total epiblast.

**Figure 2.**
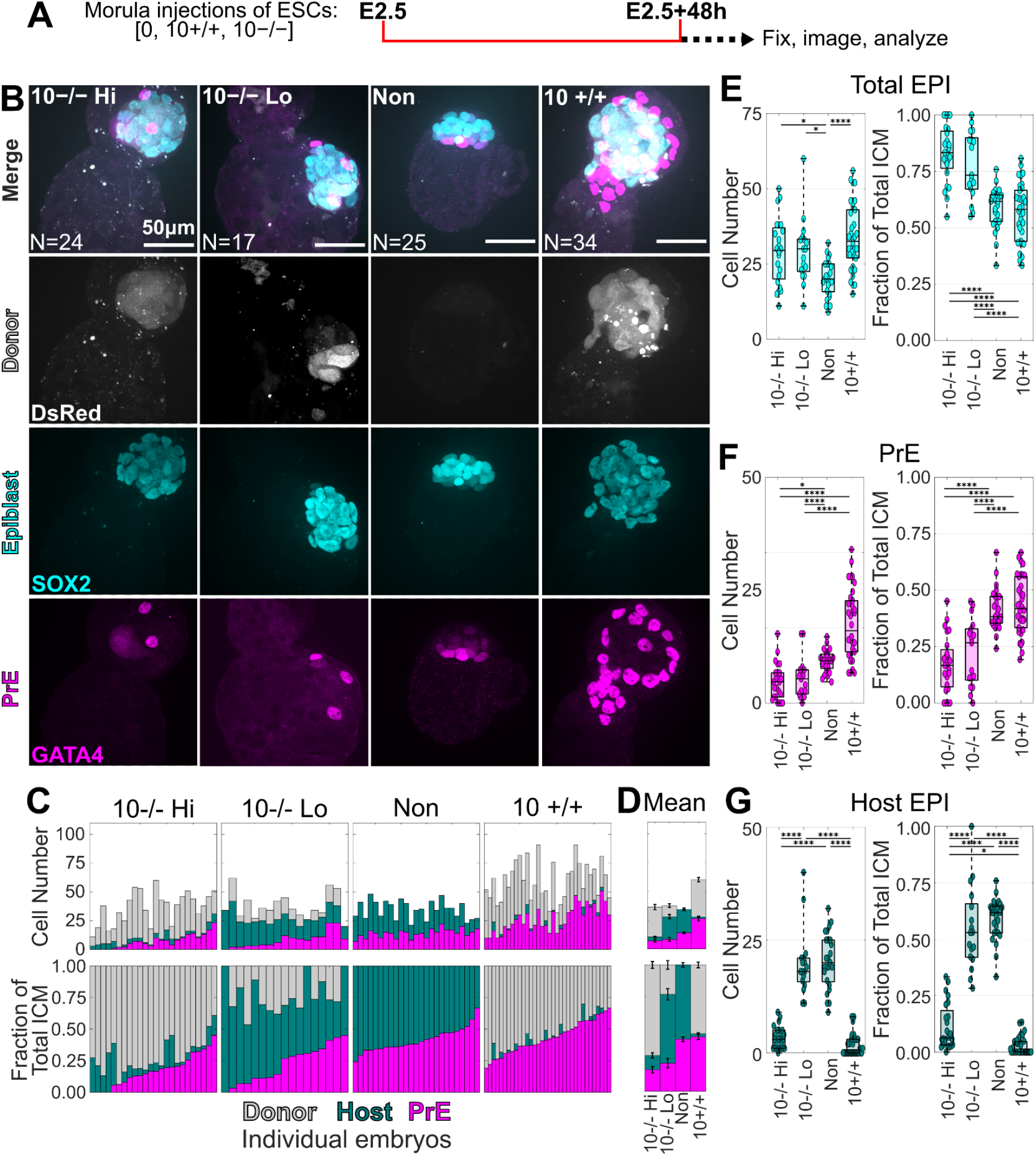
Both *Fgf4*^*+/+*^ and *Fgf4-/-* donor ESCs can impede host cells from contributing to the epiblast. **A**. Schematic for ICM imaging following injections with 10 donor cells. **B**. Representative maximum intensity projections of confocal z-stacks for 10 *Fgf4-/-* ESC injected chimeras with high (Hi) donor contribution (N=24),10 *Fgf4-/-* ESC injected chimeras with low (Lo) donor contribution (N=17), non-injected embryos (N=25), and 10 *Fgf4+/+* ESC injected chimeras (N=34). **C, D**. Stacked bar plots for individual (**C**) and mean (±SEM) (**D**) ICM composition for chimeras and embryos showing cell number (top) and cell fraction of ICM (bottom). Magenta: GATA4 positive PrE; dark green: SOX2 positive host EPI; gray: DsRed/SOX2 double-positive donor cell derived EPI. Bars are sorted by PrE fraction. **E-G**. Box / swarm plots showing cell number (left) and cell fraction of the ICM (right) for total EPI (**E**), PrE (**F**), and host derived EPI (**G**). Pairwise comparisons were carried out by N-way ANOVA, p-values: *, 0.05 ≥p > 0.01; **, 0.01 ≥ p > 0.001; ***, 0.001 ≥ p > 0.0001; ****, 0.0001 ≥ p.

All three groups of chimeras had epiblasts with greater cell numbers than non-injected embryos (**Fig. 2E**). However, when examining the epiblast fraction of the ICM, only 10-/-Hi and Lo chimeras had larger epiblast compartments. The reason for this became apparent upon inspecting the number of PrE cells within the different chimera conditions (**Fig. 2F**). Both the 10-/-Hi and 10-/-Lo chimeras showed a reduced PrE cell number and fraction of the ICM, while *Fgf4*^*+/+*^ ESC injected (10+/+) chimeras showed a reciprocal increase in both PrE cell number and fraction. Finally, 10+/+ and 10-/-Hi chimeras had fewer host cells in their epiblast as compared to the 10-/-Lo chimeras and non-injected embryos (**Fig. 2G**).

This suggests alternative mechanisms of host EPI exclusion for the *Fgf4*^*+/+*^ amd *Fgf4*^*-/-*^ donor cells. For the 10+/+ chimeras, we reason that FGF4+/+ donor ESCs are driving host ICM cells into the PrE compartment through continuous FGF4 signalling, thus precluding almost all host cells from the EPI compartment. However, the FGF4-/-donor cells obviously do not produce FGF4 ligand, therefore host epiblast exclusion must be occurring through a different mechanism. Are the cells that would have become host EPI simply lost, due to something like reduced cell-cycling or apoptosis, or, like the 10+/+ chimeras, are those cells being displaced into another tissue compartment, e.g. the TE? We address this by quantifying cell numbers and types of three lineages of the late blastocyst.

### Embryos injected with *Fgf4*^*-/-*^ donor ESCs have fewer PrE and more TE cells

We investigated how *Fgf4*^*-/-*^ donor ESCs perturb host cell allocation by quantifying cell numbers of all three lineages at the late blastocyst stage (**Fig. 3A**). We injected 10 or 15 *Fgf4*^*+/+*^ (15+/+) or *Fgf4*^*-/-*^ (15-/-) ESCs into WT 8-cell stage morulae. Chimeric and non-injected WT embryos were cultured for 48 hours to the late blastocyst stage, then fixed, stained for DAPI (nuclei), SOX2 (EPI), and GATA4 (PrE), and imaged (**Fig. 3B**). Cells were segmented using MINs with *post hoc* manual curation in ImageJ (**Fig. 3C-F**). Cell types were determined by k-means clustering based on SOX2 intensity, GATA4 intensity, and distance from the EPI centre of mass (CM), with EPI cells showing high SOX2, PrE cells high GATA4, and TE cells low intensities of both markers and greater EPI CM distance.

**Figure 3.**
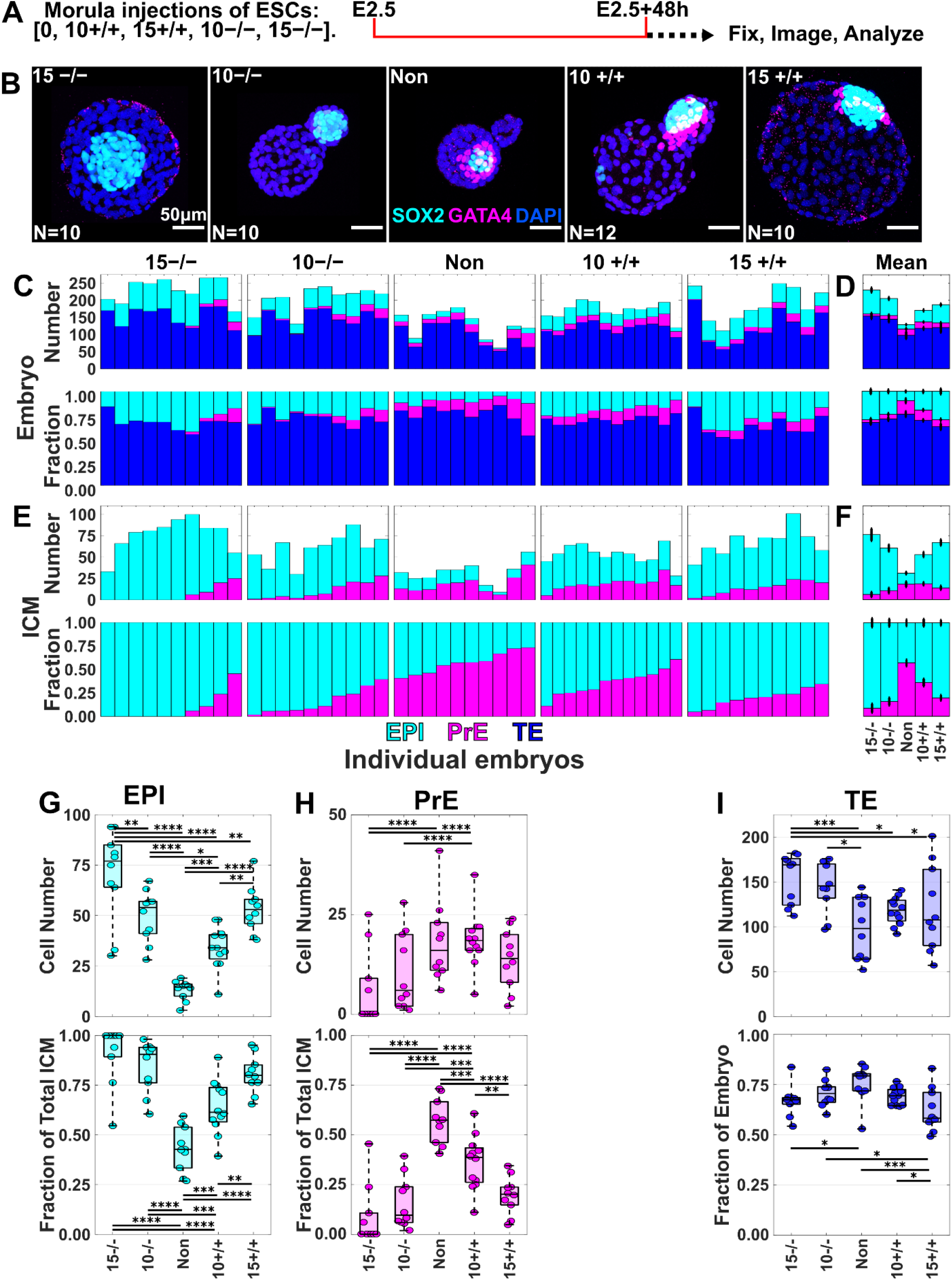
Embryos injected with *Fgf4*^*-/-*^ donor ESCs have fewer PrE cells and more TE cells. **A**. Experimental schematic for whole-embryo imaging following injections with 10 or 15 donor cells. **B**. Representative maximum intensity projections of confocal z-stacks for non-injected embryos (N=10), 10 and 15 *Fgf4*^*-/-* ESC injected chimeras (N=10,10), and 10 and 15 *Fgf4+/+* ESCl injected chimeras (N=12,10). **C, D**. Stacked bar plots for individual (**C**) and^ mean (±SEM) (**D**) ICM composition for chimeras and embryos showing cell number (top) and cell fraction of the whole-embryo (bottom). Colours show GATA4 positive PrE in magenta, SOX2 positive EPI in cyan, and double-negative TE in blue. Bars are sorted by fraction of PrE. **E**,**F**. Stacked bar plots for individual (**E**) and mean (±SEM) (**F**) ICM composition for chimeras and embryos showing cell number (top) and cell fraction of the ICM (bottom). **G**,**H**. Box plots with swarm plots showing cell number (top) and cell fraction of the ICM (bottom) for EPI (**E**) and PrE (**F**) and host derived EPI (**G**). **I**. Box / swarm plots showing cell number (top) and cell fraction of the whole-embryo (bottom) for TE. Pairwise comparisons were carried out by N-way ANOVA, p-values: *, 0.05 ≥p > 0.01; **, 0.01 ≥ p > 0.001; ***, 0.001 ≥ p > 0.0001; ****, 0.0001 ≥ p.

Consistent with the previous experiment, we observed a larger epiblast in all chimera conditions in both cell number and fraction of the ICM, as compared to non-injected embryos (**Fig. 3G**). Indeed, there was a proportional trend between the number of donor cells, of either type, and EPI number and fraction. Moreover, we saw a reduction in PrE cell number and ICM fraction for 10-/- and 15-/-chimeras (**Fig. 3H**), confirming the previous finding that *Fgf4*^*-/-*^ donor ESCs were able to impede PrE specification. In 10+/+ chimeras, we saw a numerical increase in PrE cell number, but this increase was not significant. As compared to the previous 10+/+ chimeras, we saw a decrease in the PrE fraction of the ICM in 10+/+ chimeras. The 15+/+ chimeras showed a similar trend, with an even smaller PrE fraction of the ICM. We observed that, irrespective of the donor ESC genotype, increasing the number of injected donor ESCs consistently reduced the proportion of PrE cells within the ICM.

In the *Fgf4*^*-/-*^ ESC injected embryos, the TE showed a reciprocal increase in cell number where there was a decrease in PrE cell number (**Fig. 3I**). 10+/+ chimeras showed a numerical increase in TE cell number and a subset of the 15+/+ cells have more TE cells relative to the non-injected embryos, however, neither are significant. Together this suggests that donor cells are able to drive host cells into the TE compartment during the first cell-fate decision. In the instance where the donor cells are *Fgf4*^*-/-*^, host cells specify into either EPI or PrE. Where in the case of *Fgf4*^*+/+*^ donor cells, the remaining host cells will be driven exclusively into the PrE compartment, most likely by FGF4 signalling. We next formalize these concepts in a mathematical model of chimera formation.

### Quantitative modelling suggests a role for displacement crowding and FGF4 signalling in host EPI exclusion

We integrated donor cells into our base model of embryogenesis to generate *in silico* chimeras and compared three models of chimera formation. The first model includes FGF4 induction from *Fgf4*^*+/+*^ donor cells, *D*^+^, (**Fig. 4A**) and both *Fgf4*^*+/+*^ and *Fgf4*^*-/-*^ donor cells, *D*^−^, have the same net growth rate as host cells (F) (**Fig. 4Bi**). The second model includes FGF4 induction and a different net growth rate, differential growth (GF) (**Fig. 4Bii**), and the third model includes FGF4 induction, differential growth, and spatial crowding,which drives host towards the TE, from both types of donor cells (GFC) (**Fig. 4C**). These models were simulated using the median posterior parameter values estimated from the base model of embryogenesis (**Fig. 1B, C**). Additional parameters for the GF and GFC models were estimated by ABC-MCMC using data from **Fig. 3**. Donor cell net growth rate, **α**_D_, was determined for the GF model and donor cell net growth rate, crowding factor, a, and feedback parameter, *n*, were estimated for the GFC model. Models were simulated using the median of the posterior parameter distribution (**Fig. 4D-F, Fig. S3**).

**Figure 4.**
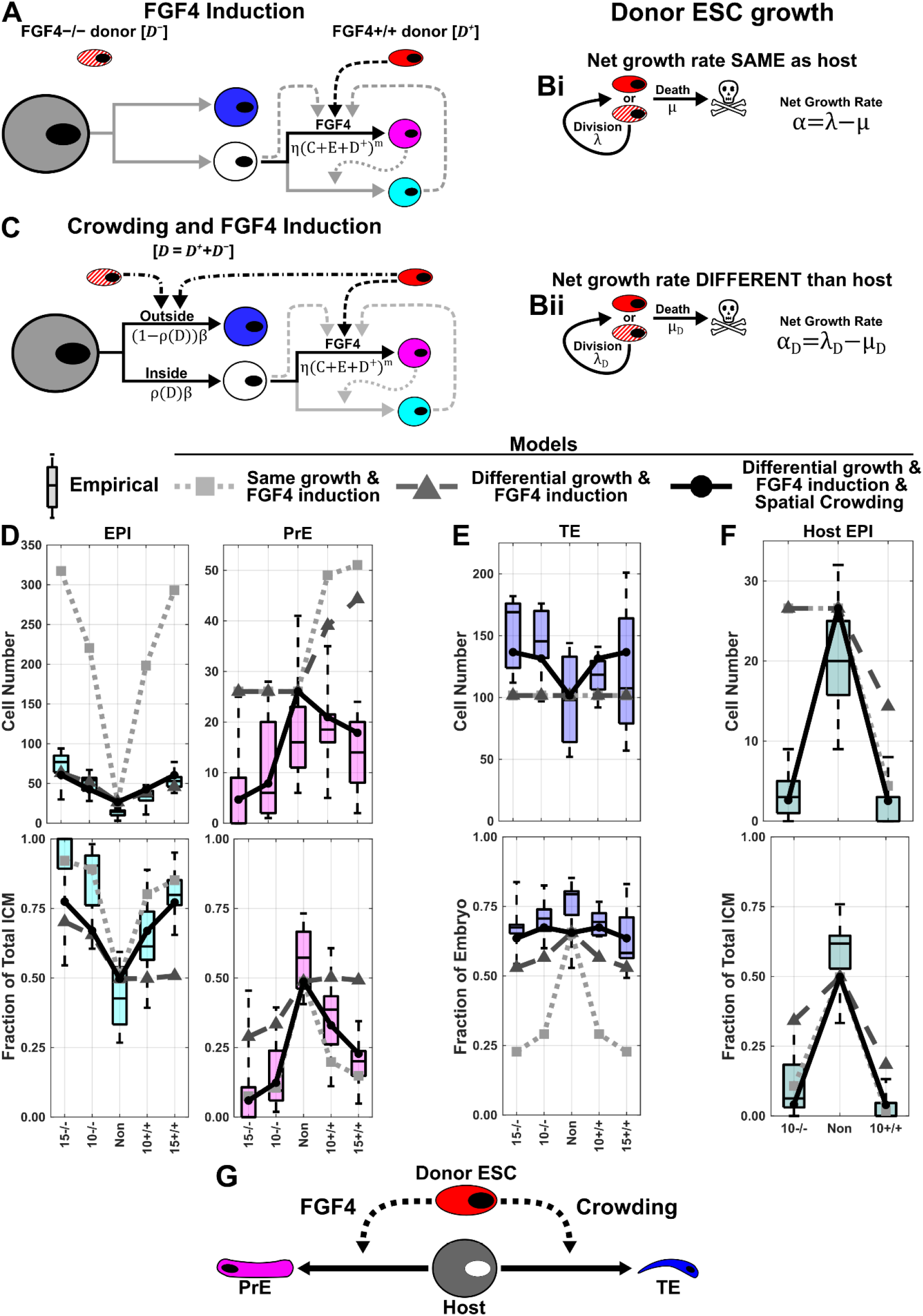
Quantitative modelling suggests a role for spatial crowding in host epiblast exclusion during chimera formation. **A**. FGF4 induction model for chimera formation. Gray: blastomere; blue: TE; white: ICM; cyan: EPI; magenta: PrE; red: *Fgf4*^*+/+*^ donor ESC; red/white: *Fgf4*^*-/-*^ donor ESC; solid lines: cell-state transitions; dashed lines: Fgf4 feedback; dotted line: posited feedback from PrE; dash-dotted line: spatial crowding feedback. **B**. Donor ESC net growth rate was either (i) fixed at the same rate as host cells or (ii) inferred. **C**. FGF4 induction and spatial crowding model for chimera formation. **D, E**. Model simulations (lines and markers) superimposed on data from **Fig. 3G-I** used to infer additional model parameters. **F**. Model simulations superimposed on host epiblast data from **Fig. 2G. G**. Proposed model for host epiblast exclusion.

Upon examining model performance, we saw the F model had EPI cell numbers more than five times larger than observed (**Fig. 4D**), while both GF and GFC models better matched EPI cell numbers. However, the GF model performed poorly against 10+/+ and 15+/+ chimeras with low EPI fractions of the ICM. Neither F nor GF models performed well against the PrE cell numbers, overestimating the values in all observed chimeras. The GFC model, on the other hand, follows the PrE trends in both cell number and fraction of the ICM. When considering the TE, the F and GF models show no modulation in TE cell number while the GFC model does (**Fig. 4E**). Finally, we test the predictive power for these models against data they have not been exposed to (**Fig. 4F**). We see that only the GFC model predicts both host EPI cell numbers and fraction of the ICM. Neither the F nor GF models are able to predict host epiblast exclusion in terms of cell number for the 10-/-chimeras. This suggests that the donor cells perturb host tissues through spatial crowding in the first cell-fate decision and FGF4 induction in the second cell-fate decision (**Fig. 4G**).

Given that ESCs are significantly smaller than the blastomeres of the E2.5 8-cell stage embryo, the ESCs may sort to the interior, while blastomeres remain on the exterior (Humięcka et al., 2016). As a result, some host cells that would otherwise be located to the interior may be forced to the exterior and specified as TE. For *Fgf4*^*-/-*^ donor ESCs, the remaining host ICM cells specify normally as either EPI or PrE, whereas for *Fgf4*^*+/+*^ donor ESCs, host cells show a bias toward PrE. Taken together, these findings advance our understanding of how donor ESC cells influence lineage allocation of host cells as ESCs reincorporate into normal development, shedding light on mechanisms of cell-fate plasticity in the early mouse embryo.

## MATERIALS AND METHODS

### ESC culture

*Fgf4*^*+/+*^ and *Fgf4*^*-/-*^ ESCs from the CD1 background (Wilder, P.J., 1997)were cultured in 2i/Lif (Ying et al., 2008) in accordance with established protocols (Mulas et al., 2019). Donor cells used in chimera experiments were labelled with dsRES via 1ug pPB-CAG-dsRED-pgk-Hyg (Guo et al., 2009), co-transfected with 2ug of transposase (pPBase; (Wang et al., 2008)), using Lipofectamine 2000. The cells were plated onto hygromycin resistant feeders and selected with 200ug/ml hygromycin after 48 hours. Individual colonies were picked after 14 days based upon dsRed expression levels.

### Embryo culture and chimaera generation

Embryos were obtained from natural mating (C57BL/6xCBA). Detection of a copulation plug in the morning was used as confirmation of successful mating and indicated embryonic day (E) 0.5. Embryos were flushed from oviducts at E2.5 8-cell stage using M2 (Sigma-Aldrich, M7167). Cells were subsequently cultured for 48 hours in BlastAssist (Origio) as either a control or following ESC injection. For chimeras, ESCs were injected via a laser-generated perforation in the zona pellucida using XYClone (Hamilton Thorne Biosciences).

This research has been regulated under the Animals (Scientific Procedures) Act 1986 Amendment Regulations 2012 following ethical review by the University of Cambridge Animal Welfare and Ethical Review Body. Use of animals in this project was approved by the ethical review committee for the University of Cambridge, and relevant Home Office licences (Project licence number 80/2597 and number P76777883) are in place.

### Immunohistochemistry

Embryos and chimeras were cultured to E2.5 + 48 hours post-harvest and fixed in 4% PFA in PBS for 15 minutes. Blocking was performed in 2% donkey serum, 0.01% BSA, 0.01% Tween20 in PBS for 15 minutes. Primary antibodies were rat monoclonal anti-Sox2 (eBioscience, 14-9811-80) at 1:500 dilution, goat polyclonal anti-Gata4 (Santa Cruz Biotechnology, SC-1237) at 1:400 dilution, and DAPI at 1:10,000 dilution. Primary antibodies were incubated at 4°C overnight. Secondary antibodies were conjugated with Alexa Fluor dyes. Confocal images were acquired using an Andor Revolution XD spinning disk confocal microscope and Leica SP5.

### Quantitative image analysis

Cell scoring and counting were performed manually on z-stacks of the embryo in Fiji using the ‘Cell Counter’ plugin. The MATLAB-based algorithm MINS (Lou et al., 2014) was used to perform a first pass of 3D nuclear segmentation of E2.5+48hrs embryos and Chimeras. Post MINS segmentation corrections for over- and under-segmentation of TE DAPI channel was performed by manual segmentation in FIJI (Schindelin et al., 2012) using ROImanager to quantify (x,y,z) coordinates and fluorescence intensity for DAPI, Sox2, and Gata4. Each segment was sorted into Sox2 single positive, Gata4 single positive, or Sox2/Gata4 double negative populations by agglomerative hierarchical clustering using MATLAB (2016B) LINKAGE with WARD distances and the CLUSTER function. Individual segments were then clustered into 3D nuclei. Optimization of the clustering and number of nuclei was performed by Scree analysis and the elbow method, respectively. Nuclei that were segmented correctly by MINS were clustered into TE, PrE, and Epiblast in the same manner as the initial segment sorting (Strawbridge et al., 2023). Clustering was performed on a per-embryo basis.

### Mathematical modelling

We modelled the mouse embryo cell population dynamics from the 8-cell stage (E2.5) to the late blastocyst stage (E4.5). Our mean-field model comprises a set of coupled ordinary differential equations (ODEs), which reflect our assumptions regarding cell state transitions, proliferation and death (**Fig. 3A, B**). These equations govern the evolution in time (*t*) of the numbers of blastomeres (*B*), TE cells (*T*), ICM cells (*C*), PrE cells (*P*), epiblast cells (*E*), and – for chimaeric embryos (**Fig. 4A-C**) – the numbers of donor ESCs that are *Fgf4*^*+/+*^ (*D*^+^) or *Fgf4*^*-/-*^ (*D*^−^).

We assumed that all host cells proliferate with constant net per-capita rate α, which captures the net effect of division and death on cell number. We modelled the sorting of blastomeres to the embryo’s interior, where they form the ICM, and exterior, where they epithelialize and become TE, as irreversible cell state transitions with constant per-capita rates ρβ and (1 − ρ)β, respectively. Here β denotes a constant overall per-capita rate of blastomere differentiation and ρ denotes the blastomere lineage bias towards the ICM. We modelled the specification of ICM into PrE and epiblast as irreversible transitions. To account for the production by PrE of extracellular matrix proteins that help drive epiblast specification, we assumed that the per-capita rate of transition from ICM to epiblast, ζ*P* ^*l*^, depends on the number of PrE cells present, with the parameter *l* allowing for non-linearity. Similarly, to account for the production by ICM and epiblast of FGF4 that helps drive PrE specification, we assumed that the per-capita rate of transition from ICM to PrE, η (*C* + *E*) ^*m*^, depends on the numbers of ICM and epiblast cells present, with the parameter *m* allowing for non-linearity. Here ζ captures the strength of feedback from PrE onto epiblast specification, while η captures the strength of feedback from ICM and epiblast onto PrE specification.

Accounting for the above processes leads to the following ODE system (**Fig. 1A**):

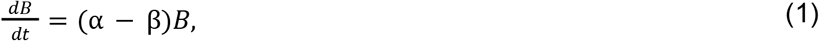

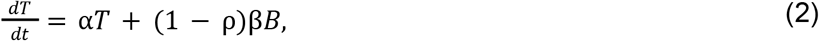

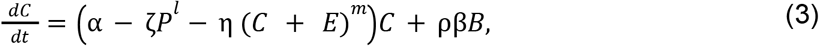

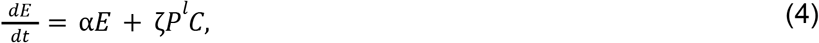

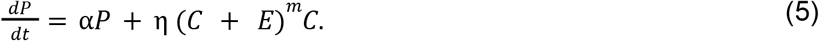

Since our model describes the cell population dynamics starting from the 8-cell stage morula comprising only blastomeres, we impose the initial conditions *B*(0) = 8, *T*(0) = *C*(0) = *E*(0) = *P*(0) = 0. Our period of interest ends at *t* = 48 hours.

Chimera formation involves the injection of donor ESCs that are *Fgf4*^*+/+*^ (*D*^+^) or *Fgf4*^*-/-*^ (*D*^−^). This introduces additional contributions to the cell population dynamics. First, we assumed that donor cells proliferate at per-capita rate, α_*D*_, which we allow to differ from that of host cells. Second, based on previous observations (Humięcka et al., 2016), we assumed that donor cells bias the rate of transition of blastomeres to ICM via crowding displacement, leading us to replace the parameter ρ with the decreasing function ρ_0_/(1 + *a*(*D* ^+^ + *D* ^−^)). We write the sum *D*^+^+ *D* ^−^ for convenience here; in practice, we only considered a model where either *Fgf4*^*+/+*^ or *Fgf4*^*-/-*^ ESCs are present. Third, to account for the production by FGF4+/+ ESCs of FGF4 that helps drive PrE specification, we modified the per-capita rate of transition from ICM to PrE from η (*C* + *E*) to η (*C* + *E* + *D*^+^). Accounting for these additional processes led to the following ODE system describing cell population dynamics during chimera formation (**Fig. 1C**):

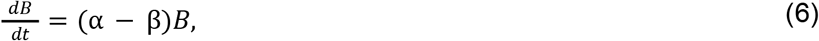

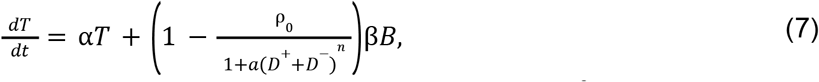

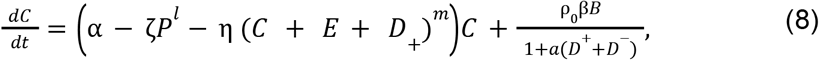

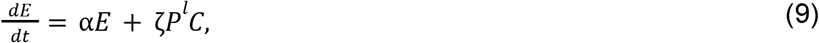

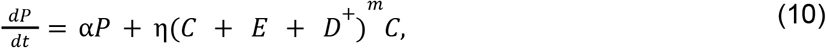

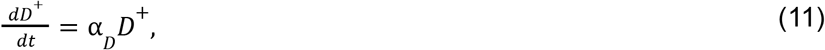

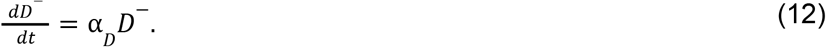

We assumed that injection of donor ESCs occurs at the 8-cell stage morula, hence imposed the initial conditions *B*(0) = 8, *T*(0) = *C*(0) = *E*(0) = *P*(0) = 0, and either *D*^+^ (0) = 10, *D*^−^ (0) = 0 (injection of 10 FGF4+/+ ESCs), *D*^+^ (0) = 15, *D*^−^ (0) = 0 (injection of 15 *Fgf4*^*+/+*^ ESCs), *D*^+^ (0) = 0, *D*^−^ (0) = 10 (injection of 10 *Fgf4*^*-/-*^ ESCs), or *D*^+^ (0) = 0, *D*^−^ (0) = 15 (injection of 15 *Fgf4*^*-/-*^ ESCs). Once again, our period of interest ends at *t* = 48 hours.

Equations (1)-(5) and (6)-(11) were solved numerically in MATLAB using an explicit Runge-Kutta method; see **Data Availability** for details on how to download our code.

Most of our model parameters cannot be directly measured and instead must be inferred from our data. We estimated parameters using Approximate Bayesian Computation (ABC) (Liepe et al., 2014; Toni et al., 2009), a likelihood-free method that iteratively compares a summary statistic from model simulations with given parameter values to the corresponding summary statistic from our data, and accepts those parameter values if these summary statistics are sufficiently close. By building up a set of accepted parameter values, ABC approximates their posterior distribution, allowing us to quantify our uncertainty in their values given our data. There are various well-established adaptations of ABC; we used a Markov Chain Monte Carlo approach (Marjoram et al., 2003). We used the summary statistic

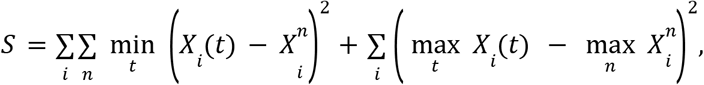

where *X*_*i*_ (*t*) denotes the value of the *i*th model variable at time *t* so that *X*_1_ (*t*) = *B*(*t*), *X*_2_ (*t*) = *C*(*t*) and so on; and 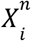 denotes the value of the *n*th observation of the *i*th variable. The first term in *S* reflects our wish to have the model solution lie as close as possible to our data, accounting for the fact that time is implicit in our experimental observations (in other words, we are fitting the model in state space rather than as a time series). The second term in *S* reflects our wish to have the maximum value attained by each component of our model solution to be as close as possible to the corresponding maximum observation, as a way of helping to pin down timescales in our model.

## ACKNOWLEDGEMENTS

The authors thank Isaac Lundie-Fallon for support with collecting and imaging embryos, Dr Néstor Saiz for discussions of complementary projects.

## COMPETING INTERESTS

The authors declare no competing financial interests.

## AUTHOR CONTRIBUTIONS

Conceptualization: S.E.S, A.G.F, J.N.; Data curation: S.E.S, A.K.S., P.H.; Formal analysis: S.E.S, A.K.S., P.H., A.G.F; Funding acquisition: S.E.S, A.G.F., J.N.; Investigation: S.E.S, A.K.S., A.G.F, J.N.; Methodology: S.E.S, A.G.F, J.N.; Project administration: S.E.S, A.G.F, J.N.; Resources: K.A.J., J.A., A.-K.H., J.N.; Software: S.E.S, A.K.S., P.H., A.G.F; Supervision: S.E.S, A.G.F, J.N.; Validation: S.E.S, A.K.S., J.A., A.G.F, J.N.; Visualization: S.E.S; Writing – original draft: S.E.S, A.G.F, J.N.; Writing – review & editing: S.E.S, A.G.F, J.N.;

## FUNDING

S.E.S. was supported by a Company of Biologists Travelling Fellowship (DEVTF-180513) and Sir Henry Wellcome Postdoctoral Fellowship (224070/Z/21/Z). A.G.F. was supported by the Biotechnology and Biological Sciences Research Council (BB/V018647/1 and BB/R016925/1).

## DATA AVAILABILITY

MATLAB code for generating all results presented in **Figs. 1D-H, 2C-G, 3C-I, 4D-F, and S1A-B** is freely available to download from GitHub (https://github.com/stanleystrawbridge/strawbridge_et_al_2024).

## SUPPLEMENTARY MATERIAL

**Figure S1.**
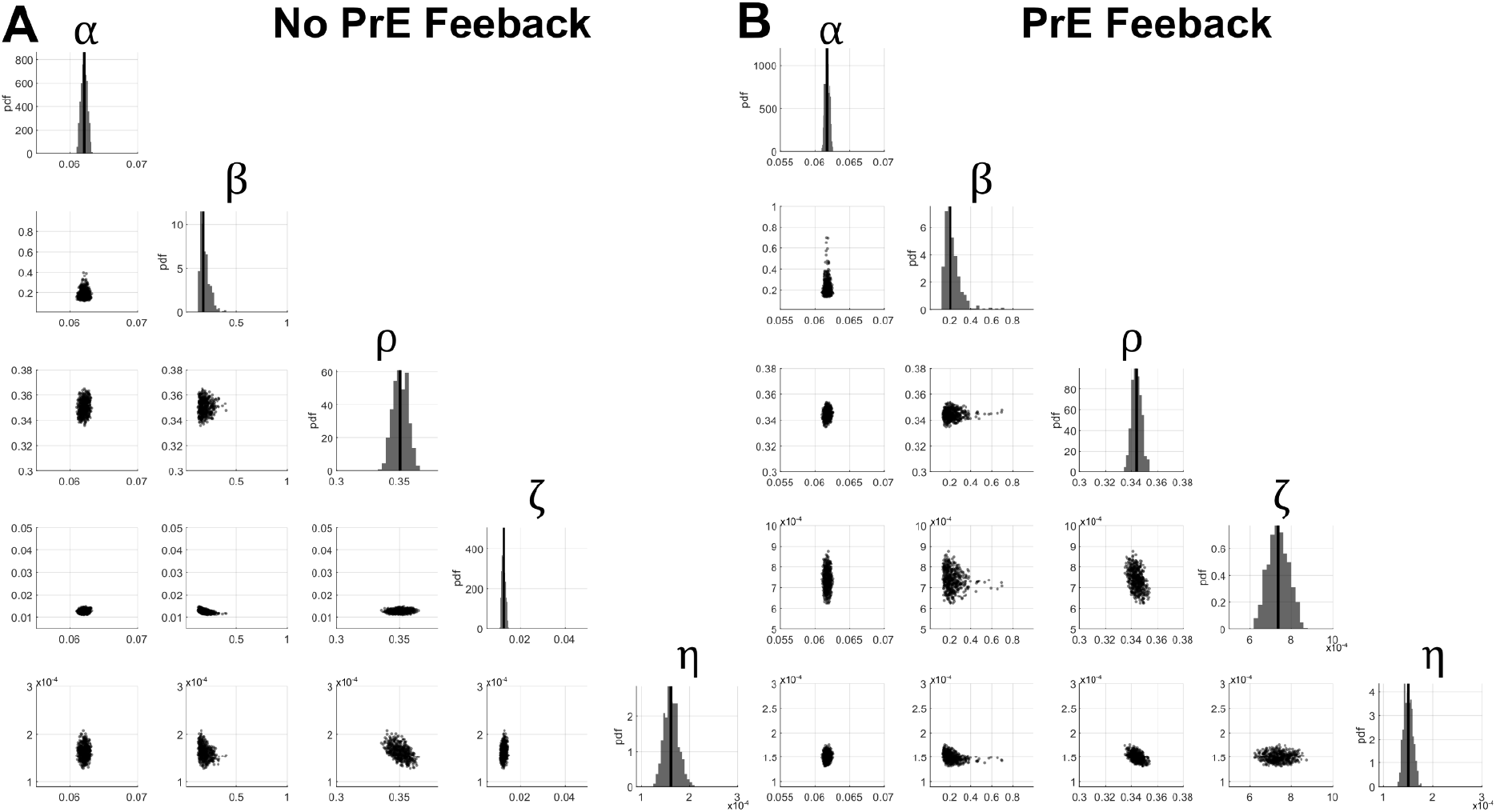
Posterior distributions of inferred parameters for different models of embryogenesis. Univariate (on diagonal histograms, vertical line shows median) and bivariate (off diagonal scatter plots) posterior distributions of inferred parameters for models without (**A**) and with (**B**) PrE feedback on ICM specification. Parameters are: **α**, net growth rate; **β**, blastomere specification rate; **ρ** blastomere bias towards the ICM fate; **ζ**, coefficient for ICM to epiblast specification; and **η**, coefficient for ICM to PrE specification. Not shown are posteriors for feedback parameters *l, m*, which both take on the value of 2 for all accepted parameter sets.

**Figure S2.**
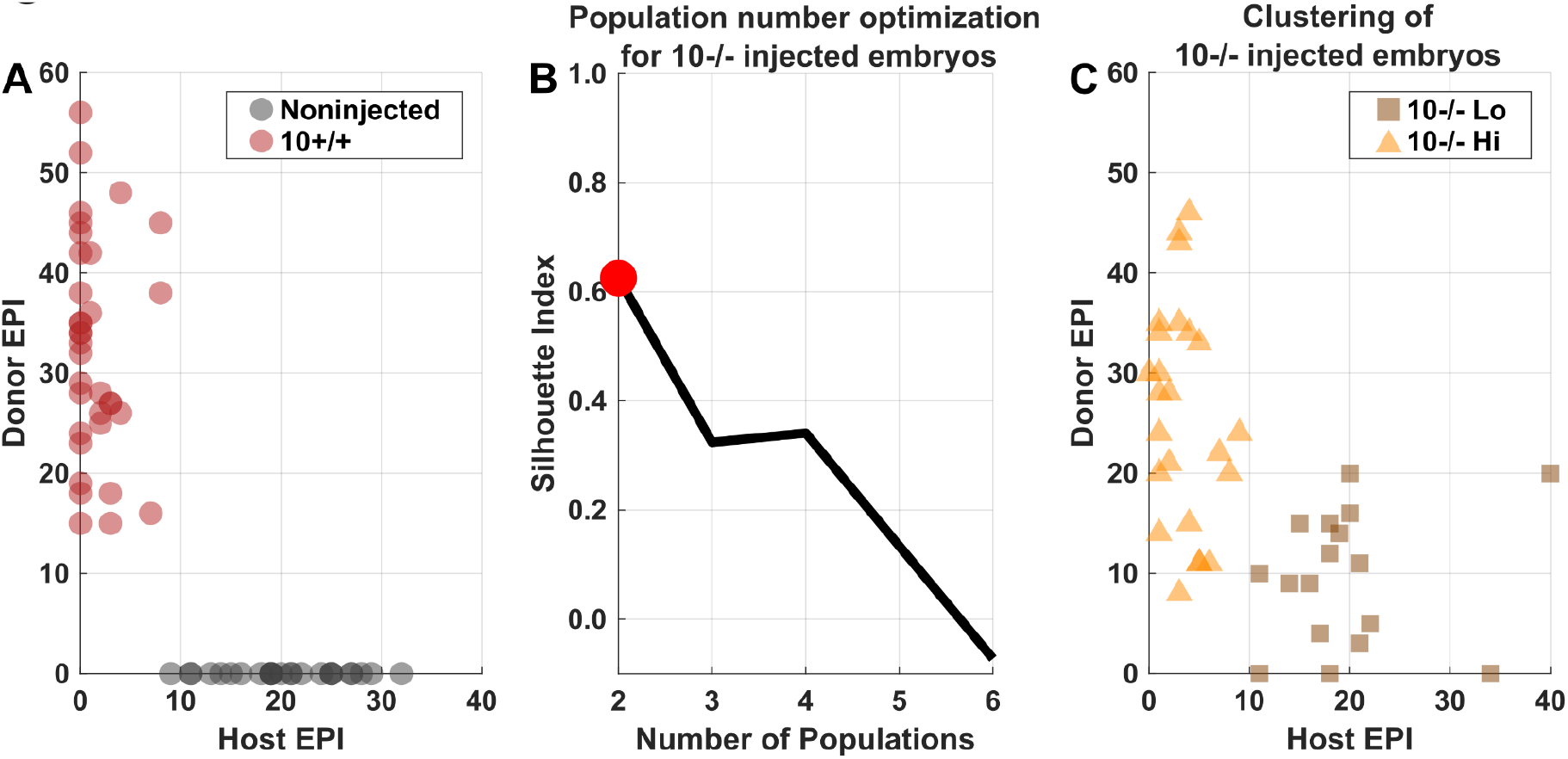
k-means clustering identifies two populations in the 10 *Fgf4*^*-/-*^ donor ESC injected embryos. **A**. Epiblast composition of noninjected (black circles) and 10 *Fgf4*^*+/+*^ donor ESC injected (red circles) embryos. **B**. Black line shows the silhouette index from k-means clustering for different numbers of populations of *Fgf4*^*+/+*^ donor ESC injected embryos. The Red dot highlights the optimal number of populations. **C**. Epiblast composition of *Fgf4*^*-/-*^ donor ESC injected embryos grouped by k-means clustering with optimal number of populations revealing a low (Lo) (brown squares) and high (Hi) (yellow triangles) donor cell contribution population.

**Figure S3.**
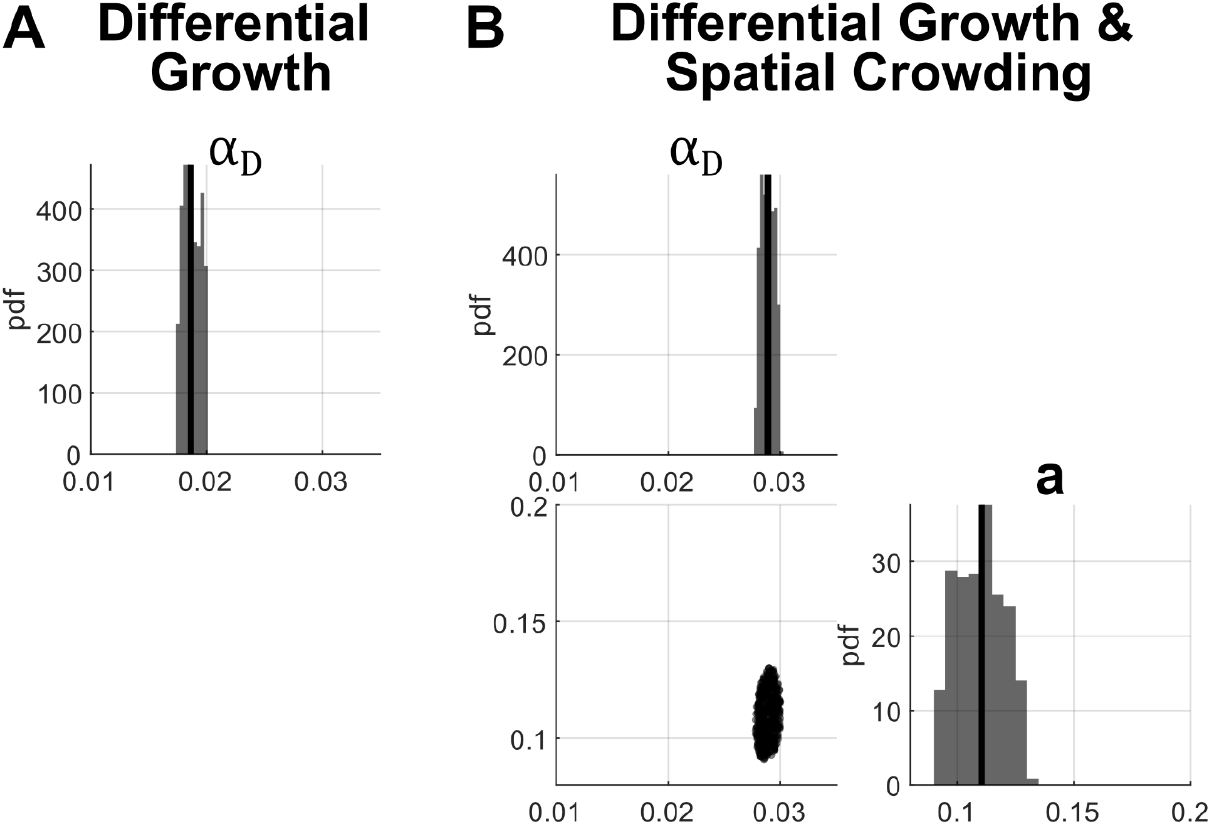
Posterior distributions of inferred parameters for different models of chimera formation. Univariate (on diagonal histograms, vertical line shows median) and bivariate (off diagonal scatter plots) posterior distributions of inferred parameters for models with differential growth for donor cells (**A**) and with differential growth and spatial crowding (**B**). Parameters are: **α**_D_, donor cell net growth rate; **a**, crowding factor. Not shown is the posterior for feedback parameter *n* which took on the value of 1 for all accepted parameter sets in the differential growth and spatial crowding model.

## Notes

### Competing Interest Statement

The authors have declared no competing interest.

